# Use of a MAIT activating ligand, 5-OP-RU, as a mucosal adjuvant in a murine model of *Vibrio cholerae* O1 vaccination

**DOI:** 10.1101/2022.06.17.496603

**Authors:** Owen Jensen, Shubhanshi Trivedi, Kelin Li, Jeffrey Aubé, J. Scott Hale, Edward T. Ryan, Daniel T. Leung

**Affiliations:** Division of Infectious Diseases, Department of Internal Medicine, University of Utah School of Medicine, Salt Lake City, USA; Division of Microbiology & Immunology, Department of Pathology, University of Utah School of Medicine, Salt Lake City, USA; Division of Chemical Biology and Medicinal Chemistry, UNC Eshelman School of Pharmacy, University of North Carolina at Chapel Hill, Chapel Hill, USA; Division of Infectious Disease, Massachusetts General Hospital, Boston, USA; Department of Medicine, Harvard Medical School, Boston, USA; Department of Immunology and Infectious diseases, Harvard School of Public Health, Boston, USA

**Keywords:** Cholera, MAIT cells, cholera vaccines, mucosal adjuvant, 5-OP-RU, conjugate vaccine

## Abstract

**Background:** Mucosal-associated invariant T (MAIT) cells are innate-like T cells enriched in the mucosa with capacity for B cell help. We hypothesize that targeting MAIT cells, using a MAIT activating ligand as an adjuvant, could improve mucosal vaccine responses to bacterial pathogens.

**Methods:** We utilized murine models of *Vibrio cholerae* vaccination to test the adjuvant potential of the MAIT activating ligand, 5-(2-oxopropylideneamino)-6-D-ribitylaminouracil (5-OP-RU). We measured *V. cholerae*-specific antibody and antibody-secreting cell responses, and used flow cytometry to examine MAIT cell and B cell phenotype, in blood, bronchoalveolar lavage fluid (BALF), and mucosal tissues, following intranasal vaccination with live *V. cholerae* O1 or a *V. cholerae* O1 polysaccharide conjugate vaccine.

**Results:** We report significant expansion of MAIT cells in the lungs of 5-OP-RU treated mice, and increases in BALF *V. cholerae* O-specific*-*polysaccharide IgG responses in our conjugate vaccine model adjuvanted with low-dose 5-OP-RU. No significant differences in humoral responses were found in our live *V. cholerae* vaccination model.

**Conclusions:** Using a murine model, we demonstrate the potential, as well as the limitations, of targeting MAIT cells to improve antibody responses to a mucosal cholera vaccine. Our study highlights the need for future research optimizing MAIT cell targeting for improving mucosal vaccines.

**One Sentence Summary:** Targeting mucosal-associated invariant T (MAIT) cells with a mucosal adjuvant in an intranasal cholera vaccine model resulted in significant expansion of lung MAIT cells, but limited improvements in cholera-specific antibody responses.

## Introduction

MAIT cells are innate-like T cells that display a semi-invariant T cell receptor (TCR) primarily composed of the TCRα chain, Vα7.2 linked to Jα33/12/20 in humans or Vα19 linked to Jα33 in mice, with a limited array of TCR β chains [1–3]. The MAIT TCR recognizes pyrimidine intermediates from the riboflavin synthesis pathway presented on the evolutionarily conserved major histocompatibility complex (MHC) Class I-related protein, MR1 [1,4–6]. MAIT cells can be activated via TCR:MR1 engagement or through cytokine stimulation, and were initially appreciated for their potent production of pro-inflammatory cytokines, including interferon-γ (IFNγ), tumor necrosis factor-α(TNFα), and interleukin-17A (IL-17A), and cytotoxic molecules [7–9]. Recent studies have shown associations between MAIT cells and adaptive immune responses, including the ability for MAIT cells to provide B cell help *in vitro* and *in vivo*, suggesting that MAIT cells might be potential targets for improving responses to mucosal vaccines, including against mucosal pathogens [10–18]. Pertinent to this, we and others have previously shown that MAIT cell frequency and activation are associated with LPS-specific antibody responses in the blood of cholera patients [10] and shigella vaccinees [11]. We have further shown that human MAIT cells are able to promote B cell differentiation and antibody production *in vitro* [12], and that T follicular helper (Tfh)-like MAIT cells in tonsils can produce key B cell help cytokines, such as IL-21 [17]. MAIT cells have also been shown to promote antibody production in a mouse model of lupus [13], a macaque model of SIV vaccination [14], and are correlated with peptide specific T cell responses in humans given adenovirus vector [16] and SARS-CoV2 mRNA vaccines [18]. Based on these data, we hypothesized that MAIT cells may be promising targets for mucosal vaccine adjuvants to improve adaptive B and T cell responses.

Cholera is a mucosal infection of global importance. Cholera is an acute dehydrating diarrheal disease caused by the non-invasive bacterial pathogen, *Vibrio cholerae*, which causes 2-5 million cases per year resulting in tens of thousands of deaths [19]. Efforts to control cholera globally include the administration of oral cholera vaccines (OCVs) (WHO, 2018). OCVs likely provide protection via the development of antibodies against *V. cholerae* lipopolysaccharide (LPS) [20]. Despite increased use in endemic areas, OCV protection remains limited with an average two-dose efficacy of 58% in adults and 30% in children under five years of age [21]. Notably, *V. cholerae* LPS memory B cell and antibody responses highly correlate with cholera protection [22,23], and these responses are markedly reduced in children under five years of age receiving OCVs compared to children with confirmed cholera infection [24]. This reduced ability to mount effective polysaccharide-specific antibody responses to vaccines is consistent with previous reports of reduced bacterial polysaccharide antibody in infants in response to polysaccharide vaccines [25]. Thus, the need for improved vaccines to enhance anti-*V. cholerae* polysaccharide immune responses in children is needed.

Mucosal adjuvants have the potential to improve mucosal vaccine efficacy through many mechanisms, including enhancing vaccine delivery, reducing tolerogenic responses, and increasing mucosal immune cell-specific targeting to improve antigen presentation and the development of adaptive immune responses [26]. In particular, using adjuvants to target innate-like lymphocytes such as invariant natural killer T (iNKT), γδ T, and mucosal-associated invariant T (MAIT) cells holds potential to improve mucosal vaccines. Targeting iNKT cells using iNKT ligands, α-galactosylceramide (α-GC) and α-GC analogs, has been studied extensively in mouse vaccination models with varying success [27]. Notably, oral α-GC gavage along with OCV in mice resulted in higher anti-LPS specific immune responses over OCV alone [28]. Despite these promising results, efficacy in humans remains uncertain as iNKT frequency in human intestines is approximately 10-fold lower than in mice [29–31]. Alternatively, MAIT cells are enriched in human blood, liver, and mucosa [7,32–36] and thus may be promising targets for OCV adjuvants.

The recent discovery of MR1 binding and MAIT-activating pyrimidine, 5-OP-RU, has allowed for improved MAIT identification via MR1-5-OP-RU tetramer binding and direct MAIT targeting *in vitro* and *in vivo* [37–44]. 5-OP-RU is highly unstable until bound by MR1 and is formed by the reaction of an early vitamin B synthesis intermediate, 5-amino-6-D-ribitylaminouracil (5-A-RU), and methylglyoxal (MGO) [37]. 5-OP-RU, when administered intranasally with toll-like receptor (TLR) agonists in mice, significantly expands MAIT cells and has been shown to improve protection in mouse models of mucosal *Salmonella, Legionella*, and *Mycobacterium* spp. challenge [38,39,41]. Recently, a more stable synthetic preparation of 5-A-RU has been developed allowing for easier storage and administration of the MAIT ligand [45]. Utilizing this tool as a mucosal vaccine adjuvant, we aimed to determine if activating MAIT cells in the presence of *V. cholerae* antigens is a viable option to improve humoral immune responses; we used live *V. cholerae* and *V. cholerae* polysaccharide conjugate antigens in murine vaccination models to address this issue. We report modest effects on mucosal immune responses in mice treated with a *V. cholerae* polysaccharide conjugate vaccine along with MAIT ligand, though no significant differences in humoral responses in our live *V. cholerae* vaccination model. Our study, by exhibiting both the potential and limitation of targeting MAIT ligands as adjuvants, adds to the growing body of work investigating MAIT ligands in the context of prophylactic and therapeutic vaccines.

## Materials and Methods

### Mice

Six to eight-week-old WT C57BL/6J female mice were acquired from the Jackson Laboratory (Bar Harbor, USA) before the start of the experiment. All mice were housed under specific pathogen free conditions and all animal experiments were performed under strict accordance with the National Institutes of Health (NIH) *Guide for Care and Use of Laboratory Animals* and institutional guidelines for animal care at the University of Utah under approved protocol no. 19-08017.

### Live *V. cholerae* O1 intranasal vaccination model

The following intranasal challenge model was adapted from Nygren *et al*. [46] and Vorkas *et al*. [41], to induce systemic and mucosal *V. cholerae*-specific antibody responses and MAIT cell expansion. In brief, lightly anesthetized mice were vaccinated intranasally on day 0 with 10^6^ colony forming units (CFU) live *V. cholerae* O1 Inaba strain N16961 with PBS, or 16.67µg Pam2CSK4 (Pam2) (Invitrogen), or 16.67µg Pam2CSK4 plus 2 mM 5-A-RU (as described in [45]) and 50 µM MGO (Sigma-Aldrich) in 50 µl (25µl per nares). On day 1, 2, and 4, mice were subsequently given either intranasal PBS or 2 mM 5-A-RU plus 50 µM MGO. All mice were boosted with 10^6^ live *V. cholerae* O1 Inaba on day 28. For 7 days following live *V. cholerae* inoculations, mice were weighed daily and monitored for signs of clinical pneumonia symptoms. Mice that lost >20% of their initial body weight were euthanized. Blood samples were collected weekly by submandibular bleeds and serum was isolated using BD microtainer SST tubes (BD Biosciences). On day 35 final blood samples were collected and then mice were euthanized using isoflurane. Bronchoalveolar lavage fluid (BALF) was collected by exposing the trachea of the euthanized mouse and, using a scalpel, cutting a small hole for catheter (23G needle inserted into polyethylene tubing) insertion. The catheter was then stabilized by tying a knot with surgical silk, and a 1 ml syringe with sterile PBS was attached to the catheter needle. PBS was then gently injected into the lungs and slowly aspirated. This process was repeated with 1 ml of PBS for a total of 2 ml of BALF collected. BALF was centrifuged at 400 g x 5 min and supernatant was collected and frozen at -80 °C for enzyme-linked immunosorbent assay (ELISA). BAL cells were resuspended in FACS buffer for flow cytometry. Lungs were then perfused with 5 ml cold PBS and single cell suspensions were prepared using the gentleMACS lung Dissociation kit (Miltenyi Biotech) according to the manufacturer’s protocol. Mediastinal lymph nodes (MLN) and spleen were then excised and ground through a 70 µm filter. Lungs and spleens were then treated with ACK lysis buffer (Thermo Fisher Scientific) to lyse red blood cells. Single cell suspensions were used for flow cytometry analysis.

### *V. cholerae* O1 O-specific-polysaccharide (OSP) vaccination model

The following *V. cholerae* OSP plus MAIT ligand vaccination model was adapted from Pankhurst *et al* [47]. On day 0, 14, and 28, mice were lightly anesthetized and vaccinated intranasally with 20 µg *V. cholerae* O1 Ogawa strain PIC158 OSP:BSA (Courtesy of the Edward Ryan lab, Harvard University, Boston, USA) and 50 µM MGO plus PBS or 75 nmol 5-A-RU in 50 µl (25 µl per nares). OSP:BSA was prepared as described [48]. Serum samples were collected as described above on day 7, 21, and 35. Mice were euthanized on day 35 and BALF, lungs and MLN were collected as described above.

### Flow cytometry

Flow cytometry was performed using standard cell surface staining techniques using directly conjugated fluorochrome antibodies and analyzed on the Cytek Aurora (Cytek Biosciences). All analysis was performed using FlowJo version 10.8.0 (BD Biosciences). Prior to surface staining, tissue single cell suspensions were incubated in Fixable Viability Dye eFluor 780 (eBioscience) to exclude dead cells, washed, and then incubated in anti-mouse CD16/CD32 Fc block (Biolegend) and unlabeled MR1-6-formylpterin-tetramer (6-FP) (NIH Tetramer core) to reduce non-specific binding. Cells were then incubated for 20 minutes at RT with mouse PE conjugated 5-OP-RU MR1-tetramer (NIH Tetramer core) and the following antibodies: anti-CD4-FITC/APC-Fire810 (clone GK1.5, Tonbo Biosciences/Biolegend), anti-CD8α-AF700 (clone 53-6.7, Biolegend), anti-TCRβ-BUV496/BV421 (clone H57-597, BD Biosciences/Biolegend), anti-CD3-BUV395 (clone 17A2, Biolegend), anti-CD69-BV510 (clone H1.2F3 Biolegend), anti-B220-PE-Cy5 (clone RA3-6B2, Biolegend), anti-CD19-BV711 (clone 6D5, Biolegend), anti-CD44-BV650 (clone IM7, Biolegend), anti-CD38-PE-Cy7 (clone 90, Biolegend), anti-IgD-BV510 (clone 11-26c.2a, Biolegend), anti-CD27-FitC (clone LG.3A10, Biolegend), anti-CD138-BV605 (clone 281-2, Biolegend).

### ELISA

Serum and BALF OSP:BSA, CT, and *V. cholerae*-specific lysate IgM, IgG and IgA ELISAs were performed as described in Jensen *et al* [17]. *V. cholerae* O1 Inaba and Ogawa specific OSP:BSA were used for the live *V. cholerae* and OSP:BSA cholera vaccination model ELISAs, respectively. For all ELISAs, serum was diluted 1:20 and BALF samples were diluted 1:2.

### Enzyme-linked immune absorbent spot (ELIspot) assay

For OSP IgG and IgA ELISpots, 96-well MultiScreen-HA filter plates (Sigma) were coated with µg/well with *V. cholerae* O1 Inaba OSP:BSA in PBS and incubated overnight at 4 °C. CT ELISpots were first coated overnight with 0.1 µg/well monosialoganglioside GM1 (Sigma) in carbonate buffer overnight at 4 °C, washed 3X with 0.05% Tween20 in PBS (PBS-T) and then incubated overnight at 4 °C with 0.25 µg/well CT (Sigma). OSP and CT ELISAs were then blocked with warm 200 µl/well of R10 media (RPMI + L-Glutamate (Gibco), 10% fetal bovine serum (FBS) + 1X Penn/Strep (Thermo) for 2 hours at 37 °C. R10 was then removed by decanting and single cell suspensions of lung and spleen cells, resuspended in R10, were added at dilutions of 1×10^5^ and 1×10^6^ cells/well and incubated at 37 °C for 5 hours. Plates were washed 3X with PBS and 3X with PBS-T and then anti-mouse IgG/IgA-biotin (Southern Biotech) diluted 1:1000 in PBS-T was added and incubated overnight at 4 °C. Plates were then washed 3X with PBS-T and incubated for 1 hour at room temperature in dark with horseradish peroxidase-Avidin conjugate (Avidin-HRP, eBioscience). Following wash, plates were developed with AEC (3 amino-9-ethyl-carbozole) and enumerated compared to negative control wells.

### Statistical analysis

All statistical comparisons were made using two-tailed Mann-Whitney *U* tests using Prism version 9.2.0 (GraphPad Software, La Jolla, CA, USA). All graphs were also created using Prism.

## Results

### MAIT cells expand and persist in lungs and bronchoalveolar lavage fluid (BALF) following intranasal 5-A-RU treatment

To study the mucosal adjuvant capacity of the MAIT activating ligand, 5-OP-RU, we utilized an intranasal prime-boost vaccination model with live *Vibrio cholerae* (*V*.*c*), previously shown by our group [17] and others [46] to induce mucosal and systemic *V*.*c*-specific LPS and CT antibody responses in wildtype (WT) C57Bl/6 (B6) mice. B6 mice were inoculated on day 0 and day 28 with 1×10^6^ CFU live *V*.*c* O1 Inaba along with either toll-like receptor (TLR) 2/6 agonist, Pam2CSK4 (Pam2) alone, or in combination with four doses of 5-A-RU and MGO as outlined in Fig. 1A. At day 35 (7 days post boost) mice were euthanized and we analyzed the frequency and activation of MAIT cells in lungs, BALF, mediastinal lymph nodes (MLN), and spleens by flow cytometry. We defined MAIT cells as live CD19^-^ CD3^+^ CD44^High^ TCRβ+ MR1-tetramer^+^ (fig. S1A) and measured MAIT cell activation based on expression of CD69. In line with previous reports, we found significantly higher MAIT cell frequency in the lungs of 5-A-RU treated mice (median=2.6%) compared to the PBS (*V*.*c* only) (median=0.08%), and Pam2 (*V*.*c* + Pam2) (median=0.15%) groups (Fig. 1B & C). MAIT cell frequency was also higher in BALF (Fig. 1D), while only moderately higher in the MLN and spleen (Fig. 1E-F). Frequencies of non-MAIT CD8 and CD4 T cells were also higher in the lungs and BALF of 5-A-RU treated mice compared to the PBS group, though this is likely largely driven by TLR2/6 stimulation as the 5-A-RU group was not statistically different than the Pam2 group alone (fig. S1. B-E). Furthermore, despite higher MAIT cell frequency in lungs and BALF, we found no differences in CD69 expression frequency in any tissue at experiment endpoint (Fig 1G-J).

**Figure 1.**
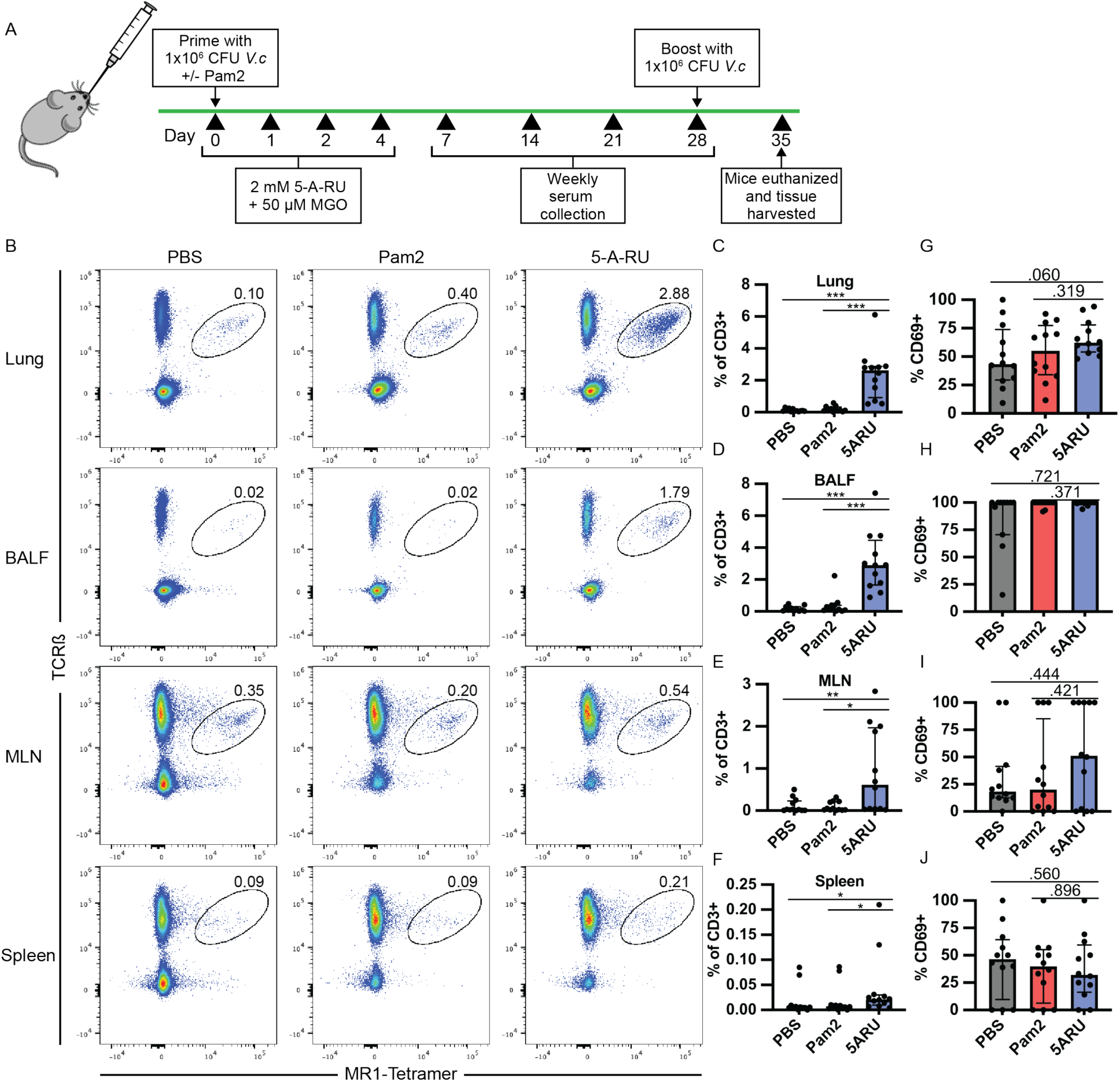
MAIT cells expand and persist in lungs and bronchoalveolar lavage fluid (BALF) following intranasal 5-A-RU treatment. (**A**) Live intranasal MAIT ligand plus *V. cholerae* O1 Inaba vaccination model timeline. (**B**) Representative FACS plots and (**C-F**) frequency as a percentage of CD3^+^ cells of MAIT cells (TCR +MR1-Tetramer^+^) from (**C**) lung, (**D**) BALF, (**E**) MLN, and (**F**) spleen gated off of Live CD3^+^ CD19^-^ CD44^+^ cells. (**G-J**) Frequency of MAIT cells expressing CD69 in (**G**) lung, (**H**) BALF, (**I**) MLN, and (**J**) spleen. Data are represented as Median with IQR from 3 independent experiments. n=11-12 mice per group. *p < 0.05, **p > 0.01, ***p > 0.001, ***p > 0.0001 by two-tailed Mann-Whitney *U* test.

### Intranasal 5-A-RU has no effect on protein or polysaccharide-specific humoral responses

Given that our previous demonstration of the sufficiency of MAIT cells to promote *V. cholerae*-specific antibody responses, and the associations of MAIT cells with LPS-specific antibody responses following human *V. cholerae* O1 infection [10] and *Shigella* vaccination [11], we hypothesized that MAIT expansion in the mouse mucosa may lead to increased LPS antibody responses. To this end, we collected weekly serum samples and BALF at endpoint and measured IgG, IgA, and IgM antibody responses against CT (protein antigen), *V*.*c* LPS O-specific polysaccharide (OSP) (polysaccharide antigen), and total *V*.*c* lysate (Fig. 2 & fig. S2). CT and OSP are the immunodominant antigens following *V. cholerae* O1 infection [20], and OSP responses are associated with protection against subsequent *V. cholerae* O1 infection [49,50]. In concurrence with our previous work [17], the PBS adjuvanted group (receiving only *V*.*c*) induced robust serum CT-IgG, CT-IgA, and OSP-IgG responses, though variable OSP-IgA responses (Fig. 2A-D). In addition to systemic antibody responses, we also recorded strong mucosal BALF CT and OSP antibody responses (Fig. 2E-H). We did not find any significant differences between either the Pam2 or the 5-A-RU group, compared to the PBS group, in any systemic or mucosal CT-or OSP-specific antibody responses at any days. We found non-significantly higher BALF OSP-IgG (P=0.212) and IgA (P=0.200) in the 5-A-RU groups (Fig. 2G & H). Additionally, no statistically significant differences were found in any systemic or mucosal CT-IgM, OSP-IgM or *V*.*c*-lysate-specific antibody responses as determined by ELISA (fig. S2).

**Figure 2.**
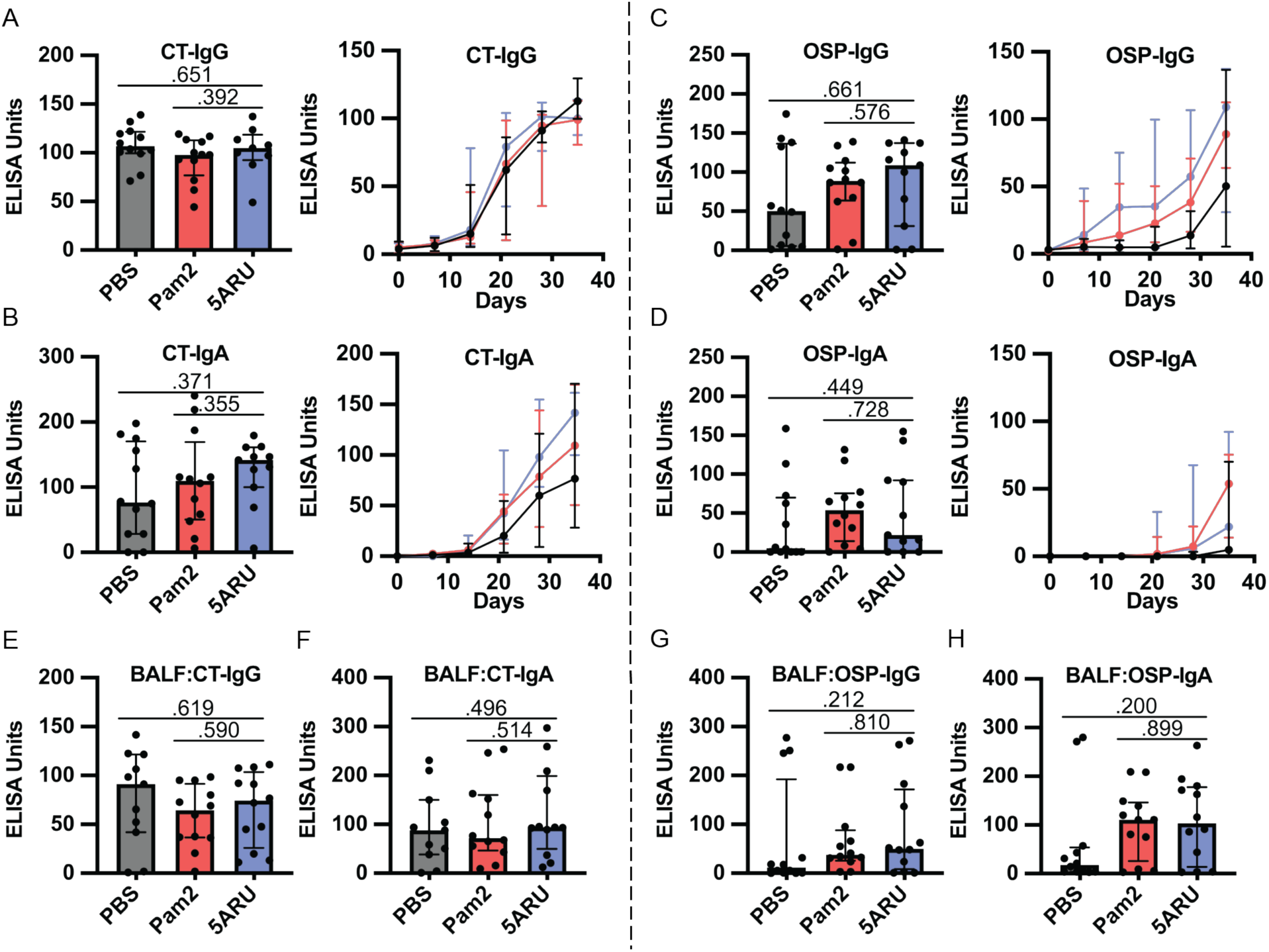
Intranasal 5-A-RU has no effect on protein or polysaccharide-specific antibody responses when administered with a live *V. cholerae* vaccination. (**A-D**) Serum day 35 endpoint (left) and time-course (right)**A**)CT-IgG, (**B**) CT-IgA, (**C**) OSP-IgG, and (**D**) OSP-IgA ELISAs. (**E-H**) BAL fluid day 35 endpoint (**E**) CT-IgG, (**F**) CT-IgA, (**G**) OSP-IgG, and (**H**) OSP-IgA ELISAs. Data are represented as ELISA units measured kinetically and normalized to positive control pooled serum from WT B6 mice intranasally vaccinated with live *V. cholerae*. Data are represented as Median with IQR from 3 independent experiments. n=11-12 mice per group. *p < 0.05, **p > 0.01, ***p > 0.001, ***p > 0.0001 by two-tailed Mann-Whitney *U* test.

In addition to antibody responses, we also examined the development of IgG and IgA OSP and CT specific antibody secreting cells (ASC) in both the mucosa (lungs) and secondary lymphoid organs (spleen) using ELISpots following intranasal 5-A-RU treatment. We found no significant differences in CT-IgG (Fig. 3A-C) or CT-IgA (Fig. 3A, D & E) ASCs in lungs or spleen between 5-A-RU and Pam2 or PBS groups. Despite clear OSP-IgG antibodies in both serum (Fig. 2C) and BALF (Fig. 2G), we found very few (<15 ASC per 1×10^6^ cells) OSP-IgG ASCs in both lungs and spleen of all groups (Fig. 3F-H). OSP-IgA ASCs were significantly higher in lungs of the 5-A-RU group compared to the PBS group (P=0.042), though were similar to the Pam2 group (Fig. 3F & I). Similarly, we report a non-statistically significant higher splenic OSP-IgA ASCs in the 5-A-RU group compared to the PBS group (P=0.150). Overall, despite substantial expansion and persistence of MAIT cells in both the mucosa and secondary lymphoid organs following 5-A-RU treatment, we found little difference in humoral pathogen-specific immune responses.

**Figure 3.**
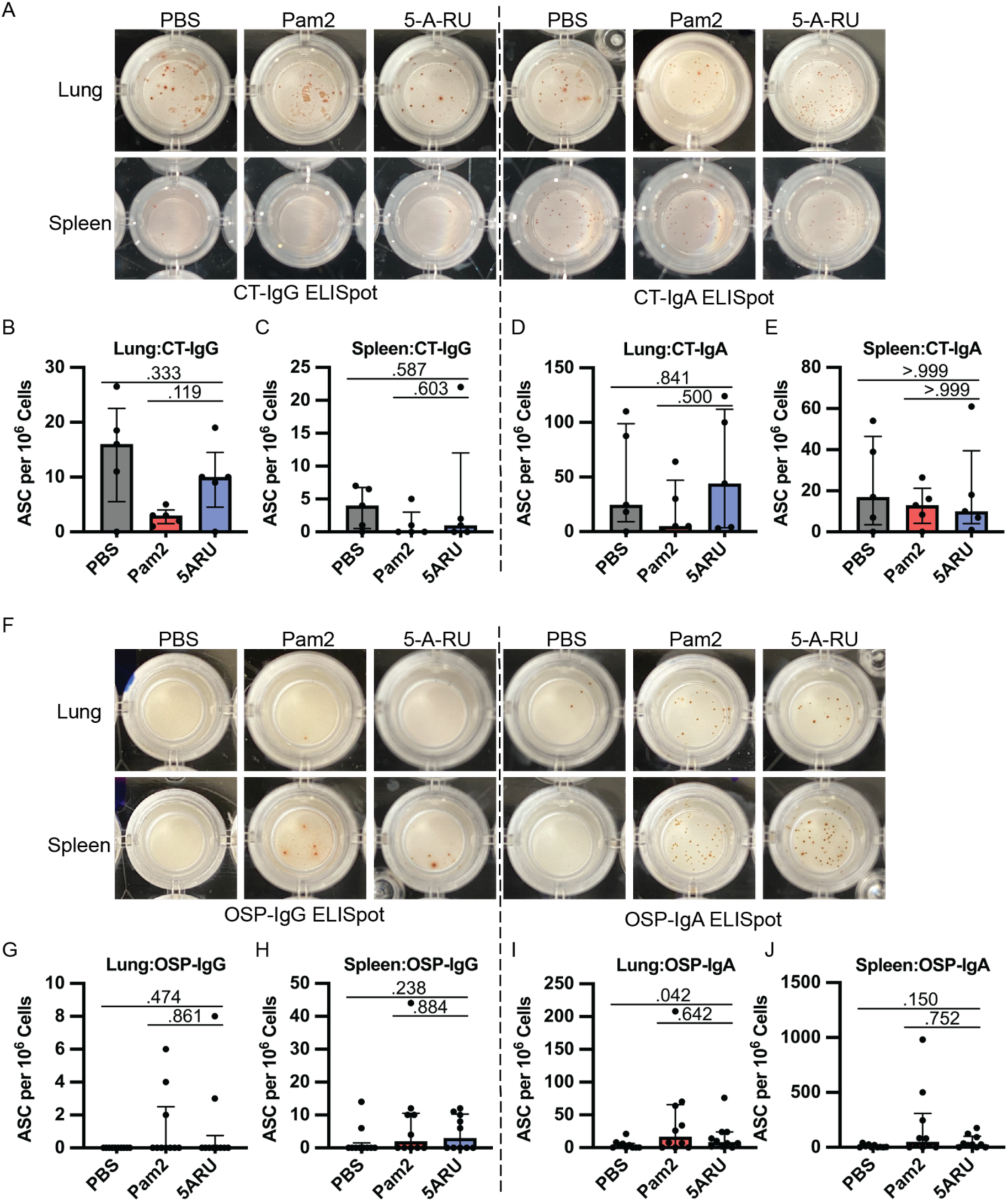
Intranasal 5-A-RU has no effect on protein or polysaccharide-specific antibody-secreting cell frequency in live *V. cholerae* vaccination. (**A**) Representative CT-IgG (left) and CT-IgA (right) ELISpot images from day 35 endpoint lung (top) and spleen (bottom) cells. (**B-E**) lung and spleen CT-IgG (**B-C**) and CT-IgA (**D-E**) ASC frequency per 10^6^ live cells. Data are represented as Median with IQR from 1 independent experiment. n=5 mice per group. (**F**) Representative OSP-IgG (left) and OSP-IgA (right) ELISpot images from day 35 endpoint lung (top) and spleen (bottom) cells. (**G-J**) lung and spleen OSP-IgG (**G-H**) and OSP-IgA (**I-J**) ASC frequency per 10^6^ live cells. Data are represented as Median with IQR from 2 independent experiments. n=10 mice per group. p values determined by two-tailed Mann-Whitney *U* test.

### MAIT cells expand following a low-dose 5-A-RU treatment combined with intranasal *V. cholerae* OSP

Due to the relatively robust anti-CT and OSP responses following the live *V*.*c* intranasal challenge (Fig. 1-3), we hypothesized that any potential adjuvant impact of intranasal 5-A-RU on *V*.*c*-specific antibody responses may be undetectable due to a ceiling effect. In support of this hypothesis, a recent study (Pankhurst et al. [47]) demonstrated that low-dose 5-A-RU augmented influenza hemagglutinin (HA) mucosal and systemic antibody responses. Thus, we sought to determine if a low-dose 5-A-RU treatment would enhance B cell responses to an intranasal vaccination of *V*.*c* O1 Ogawa OSP conjugated to bovine serum albumin (OSP:BSA). We inoculated mice on day 0, 14, and 28 with 20 µg of *V*.*c* Ogawa OSP:BSA and 100 nmol MGO +/-75 nmol 5-A-RU. Serum samples were collected on day 7 and 21, and mice were euthanized and tissue harvested on day 35 (Fig. 4A). To confirm that low dose 5-A-RU treatment without TLR agonists was effective at targeting MAIT cells, we analyzed MAIT and non-MAIT T cell frequency in the lungs, BALF, and MLN using flow cytometry. MAIT cells were defined as live B220^-^ TCRβ^+^ CD44^High^ MR1-tetramer^+^ cells (fig. S3A). We found significantly higher MAIT frequency in the lungs (P<0.001) and BALF (P<0.001) of 5-A-RU treated mice compared to MGO alone comprising a median of 6.6% and 10% of total TCRβ^+^ T cells respectively (Fig. 4B-D). In contrast, MAIT cell frequency in MLNs remained unchanged between MGO and 5-A-RU groups (Fig. 4B & E). No significant differences between groups were found in frequencies of non-MAIT CD8 and CD4 T cells in all tissues (Fig. 4C-E).

**Figure 4.**
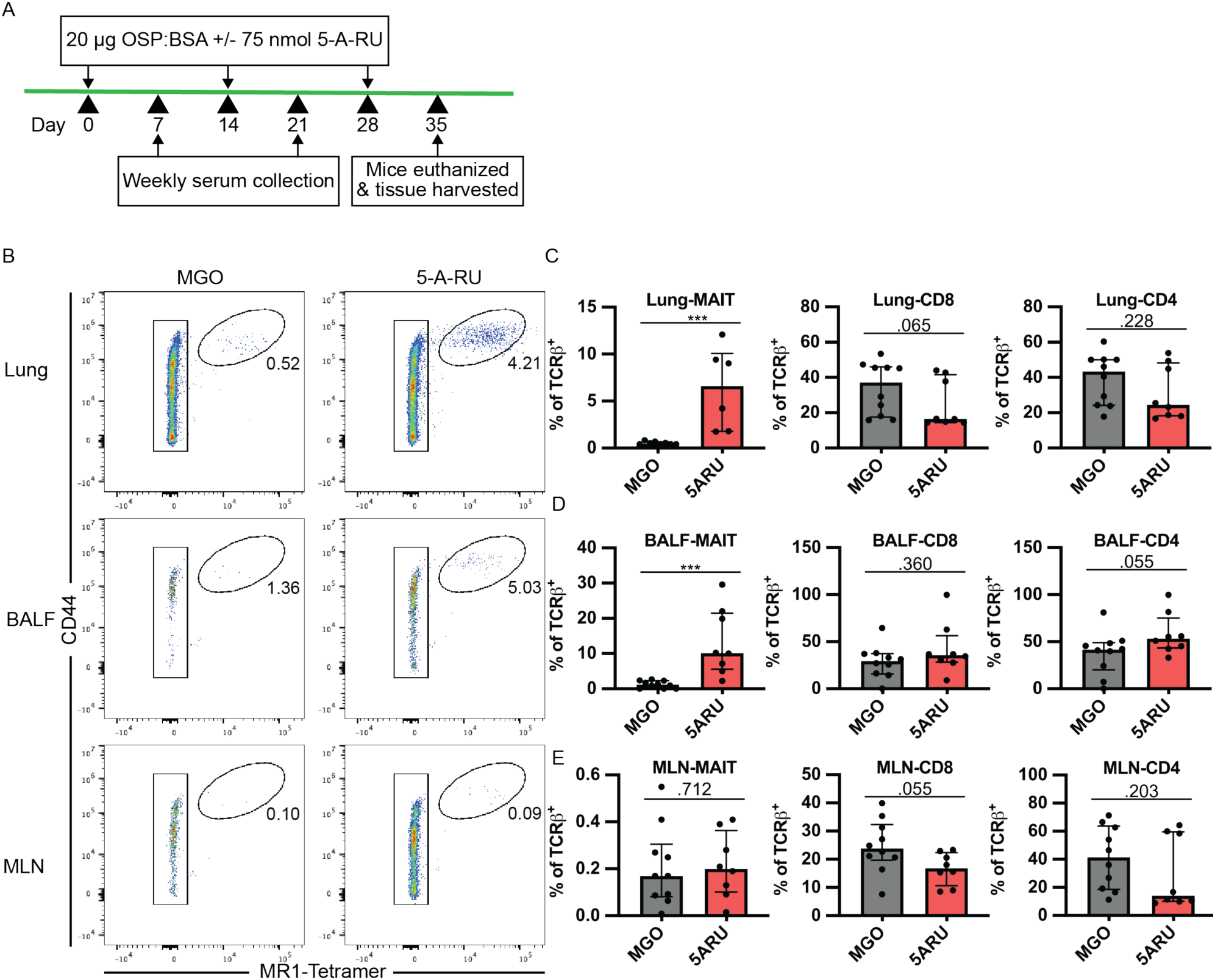
MAIT cells expand following a low-dose 5-A-RU treatment combined with intranasal *V. cholerae* OSP. (**A**) Intranasal MAIT ligand plus *V. cholerae* O1 Ogawa OSP:BSA vaccination timeline. (**B**) Representative FACS plots of MAIT cells (CD44^High^ MR1-Tetramer^+^) from lung, BALF, and MLN gated off Live B220^-^ TCRβ^+^ cells. (**C-E**) Frequency of MAIT (left), CD8 (middle), and CD4 (right) T cells as a percentage of TCRβ^+^ cells in (**C**) lung, (**D**) BALF, and (**E**) MLN. Data are represented as Median with IQR from 2 independent experiments. n=8-10 mice per group. *p < 0.05, **p > 0.01, ***p > 0.001, ***p > 0.0001 by two-tailed Mann-Whitney *U* test.

### Moderate increase in mucosal *V*.*c* OSP specific IgG following 5-A-RU treatment

After establishing that low dose 5-A-RU expands MAIT cells in the mucosa, we next sought to test whether co-administration of 5-A-RU with *V*.*c* OSP:BSA would enhance anti-OSP specific (polysaccharide) antibody responses. To test this hypothesis, we examined OSP-specific antibody responses in serum and BALF following OSP intranasal immunization with or without 5-A-RU adjuvant. We report no differences in serum OSP-IgM antibody responses between MGO and 5-A-RU (fig. S4A), and non-statistically significant higher OSP-IgG (P=0.210; Fig. 5A & B). Notably, we saw statistically significantly (P=0.045) higher BALF OSP-IgG antibodies in the 5-A-RU relative to the MGO alone group (Fig. 5C). Despite the mucosal nature of the infection, serum and BALF OSP-IgA (fig. S4C & D) and OSP-IgM (fig. S4A & B) responses in almost all mice were all below the ELISA LOD (dotted line). To investigate the relationship between OSP antibody response and MAIT cell frequency in BALF and lungs (Fig. 4B-D) we used simple linear regression analysis. Analyzing only 5-A-RU treated mice, we found statistically significant associations between BALF MAIT frequency and BALF OSP-IgG antibody responses(P=0.0389) and lung MAIT frequency and serum OSP-IgM antibody responses (P=0.0287) (Table S1). In addition to measuring antibody responses, we also used flow cytometry to analyze B cell differentiation and class switching in lung, BALF, and MLN. Specifically, we analyzed the frequencies of total B220^+^ B cells (Fig. 5G-I), IgD^+^ naïve B cells (Fig. 5J-L), and IgD^-^ CD38^high^ class switched memory B cells (Fig. 5M-O) in MGO compared to 5-A-RU groups. We found no statistically significant changes in B cell subtypes in all tissues.

**Figure 5.**
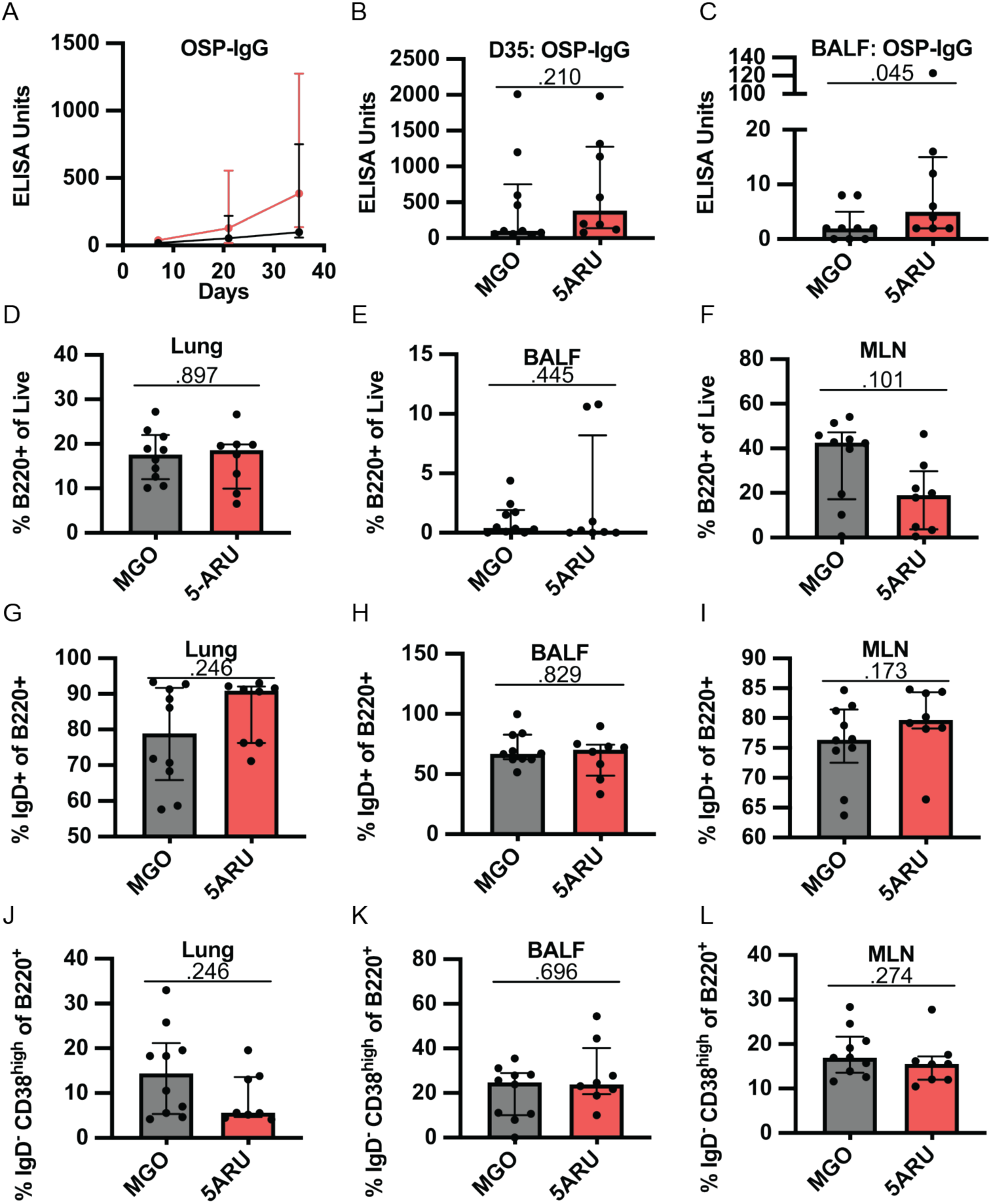
Intranasal 5-A-RU administration results in moderately higher mucosal polysaccharide responses. (**A**) Serum time-course, (**B**) serum day 35 endpoint, and (**C**) BALF day 35 endpoint OSP-IgG ELISAs. Data are represented as ELISA units measured kinetically and normalized to positive control pooled serum from WT B6 mice intranasally vaccinated with live *V. cholerae*. (**D-F**) Frequency of total B cells (B220^+^) as a percentage of live cells in (**D**) lung, (**E**) BALF, and (**F**) MLN. (**G-I**) Frequency of naive B cells (IgD^+^) as a percentage of B220^+^ cells in (**G**) lung, (**H**) BALF, and (**I**) MLN. (**J-L**) Frequency of class-switched memory B cells (IgD^-^ CD38^High^) as a percentage of B220^+^ cells in (**J**) lung, (**K**) BALF, and (**L**) MLN. Data are represented as Median with IQR from 2 independent experiments. n=8-10 mice per group. p values determined by two-tailed Mann-Whitney *U* test.

Together, these data show that low-dose intranasal 5-A-RU with MGO can expand MAIT cells in the mucosa of mice and moderately enhance mucosal IgG polysaccharide responses despite little effect on systemic antibody and B cell differentiation.

## Discussion

The recent discovery and accessibility of MR1-binding and MAIT activating ligands has led to research into potential approaches targeting MAIT cells for both therapeutic and prophylactic treatments and vaccines against cancer [43,51,52], viruses [47], and bacteria [38–41,44,53]. Our study focuses on targeting MAIT cells in the context of cholera vaccines, which have limited long term efficacy, particularly in young children [21,54,55]. In this study, we used murine models of cholera vaccination to test the adjuvant effects of targeting MAIT cells via intranasal administration of 5-A-RU with MGO. While we demonstrated significant expansion and persistence of mucosal MAIT cells through high and low-dose intranasal 5-A-RU treatment, we saw limited 5-A-RU adjuvant effects on *V*.*c*-specific antibody responses and B cell differentiation.

Among our findings was higher mucosal OSP-IgG antibodies in low-dose 5-A-RU treated mice compared to MGO controls (Fig. 5C), accompanied by a non-statistically significantly higher serum OSP-IgG (Fig. 5A & B). Additionally, we found significant correlations with BALF MAIT cell frequency and BALF OSP-IgG antibody responses (Table S1). These results are important in the context of cholera, as OSP-IgG and IgA antibodies are associated with protection from subsequent cholera infection, and recently were shown to inhibit *V. cholerae* motility *in vitro* [49,50,56]. Furthermore, these data build on previous reports associating human MAIT cells with polysaccharide-specific B cell responses, including after Shigella *dysenteriae* vaccination [11], and following V. cholerae O1 infection, where MAIT cells were correlated with LPS but not CT specific antibody responses [10]. The mechanism of this MAIT B cell help remains unknown despite studies by our group and others implicating MAIT cells in humoral responses in mice, non-human primates, and humans [12–15,17].

MAIT cells may help B cells via contact dependent or contact independent interactions. In support of contact dependent MAIT B cell interactions, human B cells activate MAIT cells *in vitro* via MR1 leading to increased frequency of inflammatory cytokine and CD40 Ligand (CD40L) producing MAIT cells [34,57], and B cells are required for MAIT cell development in the mouse periphery [32]. Furthermore, murine MAIT cells promote B cell autoantibody production *in vitro* in a mouse model of lupus via CD40L:CD40 signaling [13]. Based on these data, there is a potential for mouse MAIT cells to directly provide help to B cells via TCR:MR1 and CD40:CD40L co-stimulation. Alternatively, MAIT cells may provide B cell help via indirect interactions. For example, human MAIT cells in tuberculosis pleural effusions [15] and tonsils [17] have been shown to produce IL-21, a key cytokine in B cell differentiation, proliferation, and class switching [58], though this has not been confirmed in mice. MAIT cells have also been implicated in dendritic cell (DC) maturation via CD40L [59] and GM-CSF [60] production, suggesting DC licensing may play a role in B cell help. This is supported by Pankhurst *et al* [47] which outlines a model of improved Tfh and germinal center responses to influenza HA antigen following CD40L mediated MAIT licensing of DCs. As we used OSP conjugated to BSA (a protein antigen) it is possible enhancement of T cell responses via MAIT cell CD40L induced DC licensing was responsible for our phenotype.

Although optimization of the timing and dosing of 5-A-RU and OSP may yield more noteworthy outcomes, we hypothesize that an OSP:MAIT-ligand conjugate may be a more viable strategy to improve polysaccharide responses. This strategy has proven successful in a similar model using the iNKT agonist, α-GC, conjugated to *Streptococcus pneumoniae* capsule polysaccharide providing protection in mice [61]. In general, conjugate vaccines have been targeted to encapsulated bacteria including *Streptococcus pneumoniae, Neisseria meningitidis*, and *Haemophilus influenzae* using bacterial capsular polysaccharides conjugated to strong protein antigens such as tetanus toxin to induce germinal center B:T cell interactions and long-lived plasma and memory B cells [62]. Similarly, *V. cholerae* O1 OSP conjugate vaccines have also been tested in mouse models, where intramuscular or intradermal injections of OSP conjugated to tetanus toxin but not OSP alone induced anti-OSP IgG but not IgA antibodies and were protective in a *V. cholerae* neonatal challenge model [48,63]. As *V. cholerae* is a non-invasive pathogen, protection is largely driven by secreted IgM and IgA antibodies against OSP, potentially through inhibition of bacterial motility [50,56], thus induction of mucosal IgM and IgA is critical. Alternatively, targeting MAIT cells using an OSP:MAIT-ligand conjugate vaccine may hold promise to improve mucosal IgA responses as MAIT cells are enriched in the mucosa and have been shown by our group to boost *V. cholerae*-specific IgA antibodies in a mouse MAIT adoptive transfer model [17].

Other than higher mucosal OSP-IgG titers in our OSP vaccination model, we did not see significant differences in any other measures of the humoral immune responses we examined. Specifically, analysis of B cell and non-MAIT T cell frequencies, and serum and mucosal antibodies, showed no significant differences between our control and 5-A-RU treated groups in either our live *V*.*c* or OSP vaccination models. As our flow-based analysis was limited to cells harvested at the experiment’s endpoint, it is possible that differences at earlier time points may not be detected, though serum antibody data from both models indicate few differences. These data underscore the need for more in-depth studies regarding the mechanism of B cell help, and the kinetics of such. Mucosal 5-OP-RU has been utilized in both mouse and rhesus macaque models as a vaccine adjuvant or therapeutic treatment with varying success. Expansion of MAIT cells using intranasal 5-OP-RU reduced lung bacterial load in models of *Legionella longbeachae* [39] and *Mycobacterium bovis* Bacillus Calmette Guerin challenge [53], and 5-OP-RU therapeutic vaccination of *Mycobacterium tuberculosis* chronically infected mice reduced bacterial load in the lung, though prophylactic vaccination had a detrimental effect due to delayed CD4^+^ T cell priming [40]. In contrast, intratracheal administration of 5-OP-RU in *M. tuberculosis* infected macaques resulted only in exhaustion of MAIT cells, with no evidence of expansion, and no impact on clinical or microbiologic outcomes [44]. In the context of these prior studies, our data highlight the potential limitations of 5-OP-RU as a bacterial mucosal vaccine adjuvant, though its use in humans remain largely unexplored.

Our study has a number of limitations. First, given the paucity of differences in polysaccharide-specific immune responses between ligand-adjuvanted and non-adjuvanted groups, we did not exhaustively explore the kinetics or the dose-dependency of the limited phenotype we observed in this model. We also did not proceed with any infectious challenge studies to examine its impact on protection or bacterial burden. Second, our studies were performed in a murine model, the composition of MAIT cells in which may not approximate those in humans. For example, while mice have a clear differentiation of Th1-like and Th17-like MAIT cells [8], such a dichotomy is not evident in human MAIT cells, which possess a wide transcriptional landscape [42]. In addition, relative to humans, mice have a much higher proportion of iNKT cells in the mucosa potentially limiting the MAIT adjuvant phenotype in this tissue niche [29–31]. We also did not examine the transcriptional programming of expanded MAIT cells in our models, though previous studies have shown an increase in IL-17A producing MAIT cells following intranasal 5-OP-RU plus Pam2 [41]. Third, we report very little OSP-IgA antibodies (fig. SC & D) in the mucosa and serum despite the mucosal nature of our OSP:BSA vaccine model. This is in contrast to our live *V*.*c* challenge model (Fig. 1A) which induced robust mucosal OSP-IgA responses (Fig. 2H), and to studies showing markedly increased OSP antibodies following cholera infection compared to vaccination [50,55]. Thus, this may be a suboptimal model to investigate polysaccharide specific responses though it highlights the difficulties of mucosal vaccine development particularly to non-protein antigens, and need for mucosal adjuvants to enhance these responses.

As cholera and other mucosal bacterial infections remain a significant public health burden, the need for improved mucosal vaccine strategies is vital. We sought to investigate whether targeting MAIT cells is a viable option to improve antibody responses to *V. cholerae* vaccinations. Despite clear MAIT cell expansion following MAIT ligand treatment we found limited effects on adaptive immune responses. In conclusion, our study highlights both the potential and limitations of targeting MAIT cells with mucosal adjuvants to improve bacterial vaccines, and adds to the growing body of work investigating MAIT ligands in both therapeutic and prophylactic vaccines.

## Acknowledgements

We would like to thank the staff of the University of Utah Office of Comparative Medicine and Flow Cytometry Cores.

