## Supplemental Figures and Tables for "Use of a MAIT activating ligand, 5-OP-RU, as a mucosal adjuvant in a murine model of *Vibrio cholerae* O1 vaccination"

### Supplementary Figures and Tables

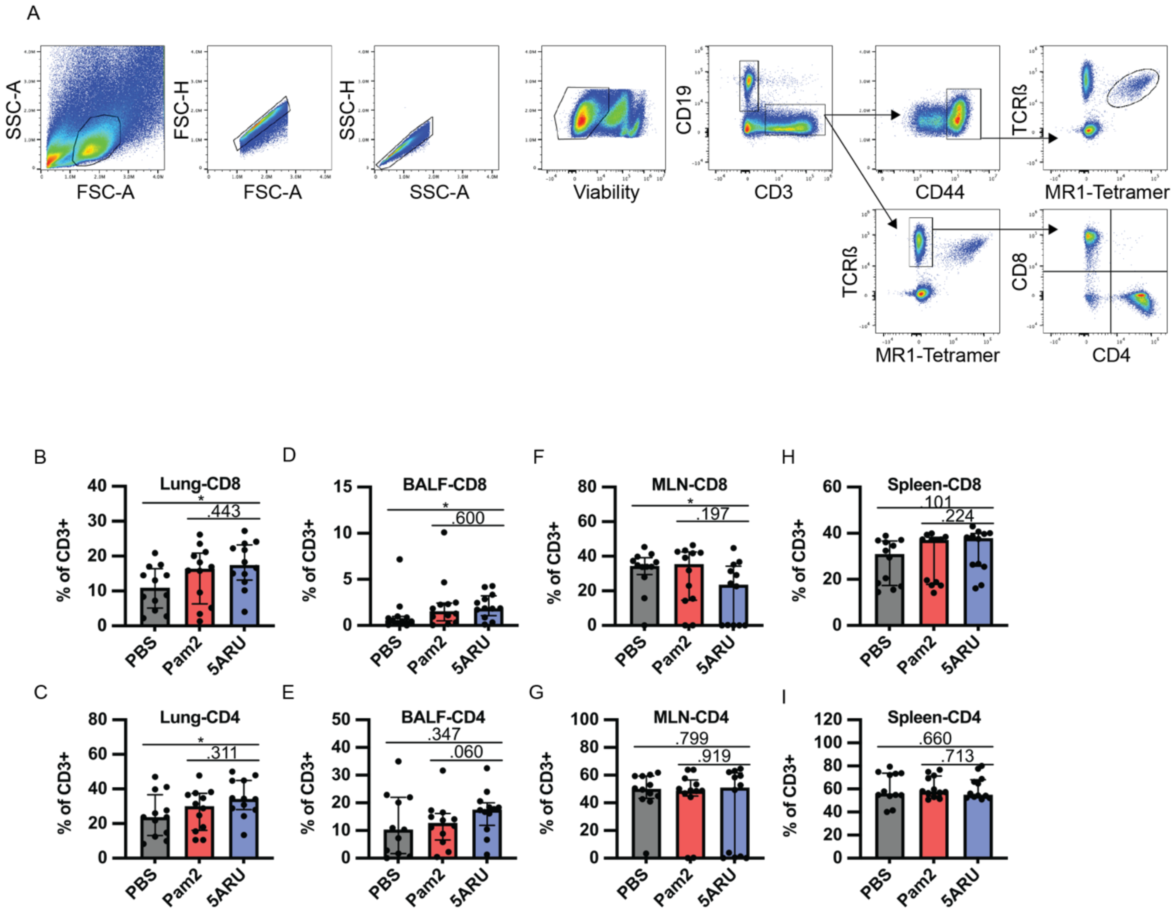


**Supplementary Figure 1. No changes in non-MAIT T cell populations in live *V. cholerae* vaccination following 5-A-RU treatment.** (**A**) Representative flow cytometry gating strategy for MAIT and non-MAIT T cells in mouse lungs vaccinated with live *V. cholerae*. (**B-I**) Frequency of CD8 and CD4 T cells as a percentage of total $CD3^{+}$ cells in Lung (**B-C**), BALF (**D-E**), MLN (**F-G**), and spleen (**H-I**). Data are represented as Median with IQR from 3 independent experiments. n=11-12 mice per group. *p < 0.05, **p > 0.01, ***p > 0.001, ***p > 0.0001 by two-tailed Mann-Whitney *U* test.

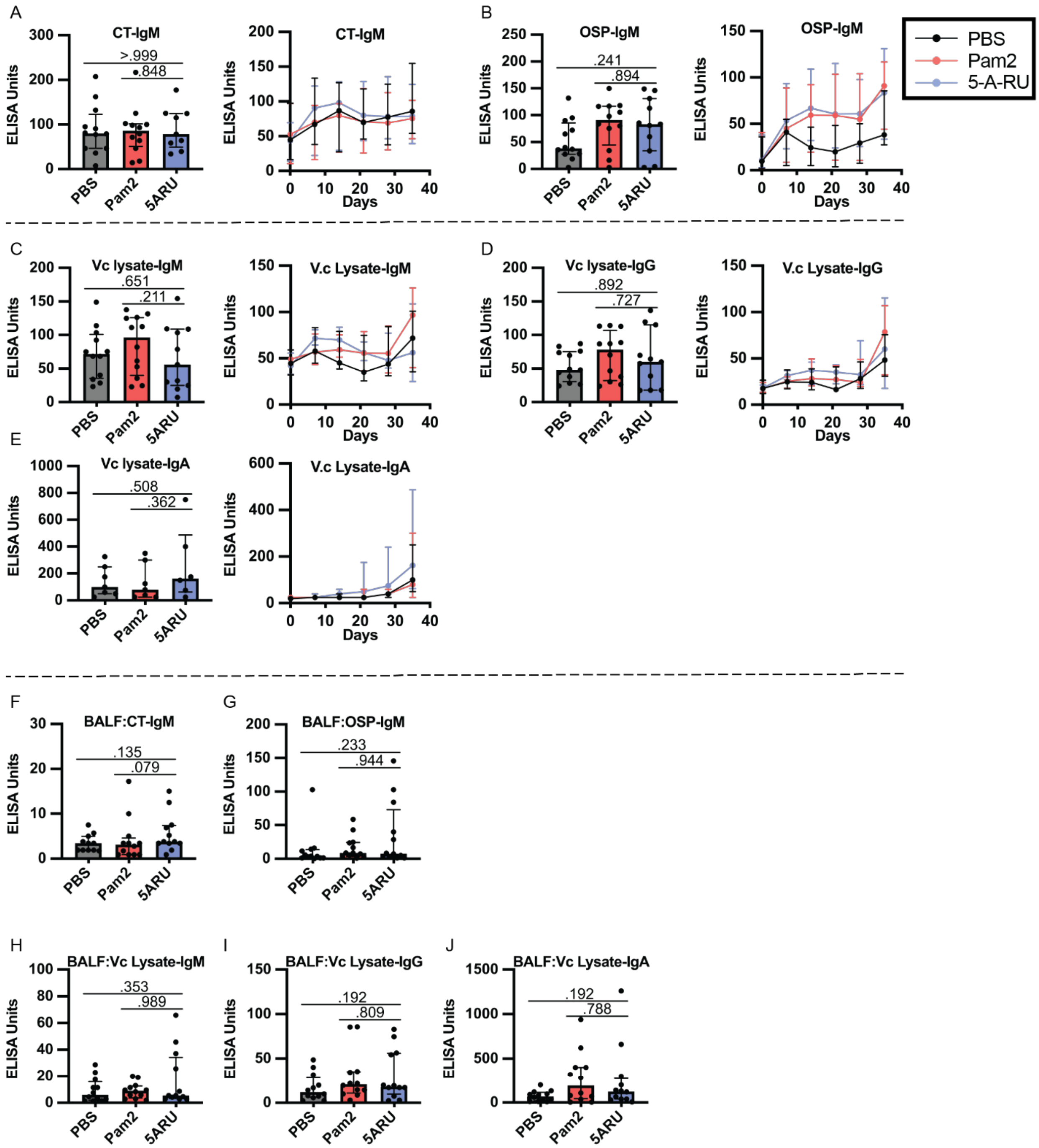


**Supplementary Figure 2. Intranasal 5-A-RU has no effect on *V. cholerae*-specific antibody responses when administered with a live *V. cholerae* vaccination.** ) Serum day 35 endpoint (left) and time-course (right) (**A**) CT-IgM, (**B**) OSP-IgM, (**C**) *V. cholerae*-lysate-IgM, (**D**) *V. cholerae*-lysate-IgG, (**E**) *V. cholerae*-lysate-IgA ELISAs. (**F-J**) BAL fluid day 35 endpoint (**F**) CT-IgM, (**G**) OSP-IgM, (**H**) *V. cholerae*-lysate-IgM, (**I**) *V. cholerae*-lysate-IgG, and (**J**) *V. cholerae*-lysate-IgA ELISAs. Data are represented as ELISA units measured kinetically and normalized to positive control pooled serum from WT B6 mice intranasally vaccinated with live *V. cholerae*. Data are represented as Median with IQR from 3 independent experiments. n=11-12 mice per group. p values determined by two-tailed Mann-Whitney *U* test.

A

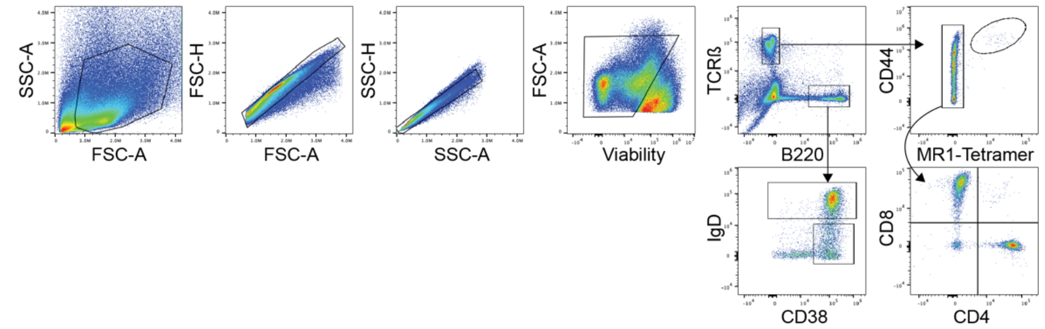


**Supplementary Figure 3. Flow gating of T and B cell populations from intranasal *V. cholerae* OSP vaccination.** (**A**) Representative flow cytometry gating strategy for MAIT cells, non-MAIT T cells and B cells in mouse lungs vaccinated with *V. cholerae* O1 Ogawa OSP:BSA.

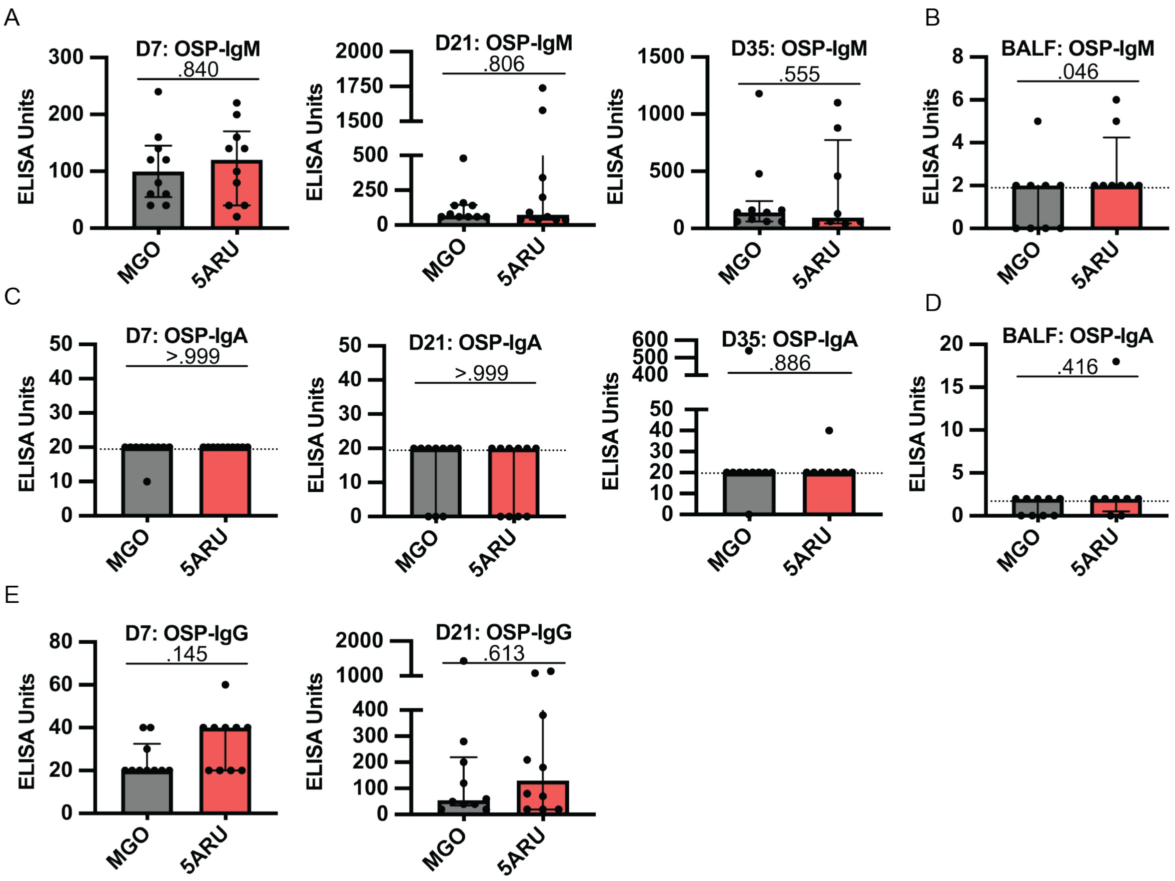


**Supplementary Figure 4. Intranasal 5-A-RU has no effect on mucosal or systemic polysaccharide-specific IgM and IgA antibody responses.** (**A**) Serum day 7 (left), day 21 (middle) and day 35 (right), and (**B**) BALF day 35 endpoint OSP-IgM ELISAs. (**C**) Serum day 7 (left), day 21 (middle) and day 35 (right), and (**D**) BALF day 35 endpoint OSP-IgA ELISAs. (**E**) Serum day 7 (left) and day 21 (right) OSP-IgG ELISAs. Data are represented as ELISA units measured kinetically and normalized to positive control pooled serum from WT B6 mice intranasally vaccinated with live *V. cholerae*. p values determined by two-tailed Mann-Whitney *U* test. Dotted lines indicate Limit of Detection (LOD).

| ELISA (Sample Tissue:Target) | MAIT Frequency (Tissue) | R squared | P Value |
|---|---|---|---|
| BALF:OSP-IgG | BALF | 0.536 | 0.0389 |
| BALF:OSP-IgG | Lung | 0.1564 | 0.4377 |
| Serum:OSP-IgG | BALF | 0.2851 | 0.1728 |
| Serum:OSP-IgG | Lung | 0.01737 | 0.8327 |
| Serum:OSP-IgM | BALF | 0.03489 | 0.2169 |
| Serum:OSP-IgM | Lung | 0.7369 | 0.0287 |

**Supplementary Table 1. OSP ELISA vs MAIT frequency.** Simple linear regression analysis associating BALF and serum OSP:BSA IgG and IgA ELISA data and MAIT frequency from BALF and lungs in mice treated with *V. cholereae* OSP:BSA and 5-A-RU (Figures 4 and 5). BALF OSP-IgM ELISA data was omitted from analysis as few samples were above the LOD.
